# The relationship between age, neural differentiation, and memory performance

**DOI:** 10.1101/345181

**Authors:** Joshua D. Koen, Nedra Hauck, Michael D. Rugg

**Affiliations:** Department of Psychology, University of Notre Dame, IN; Center for Vital Longevity, University of Texas at Dallas, TX; School of Behavioral and Brain Sciences, University of Texas at Dallas, TX; University of Texas Southwestern Medical Center, TX

## Abstract

Healthy aging is associated with decreased neural selectivity (dedifferentiation) in category-selective cortical regions. This finding has prompted the suggestion that dedifferentiation contributes to age-related cognitive decline. Consistent with this possibility, dedifferentiation has been reported to negatively correlate with fluid intelligence in older adults. Here, we examined whether dedifferentiation is associated with performance in another cognitive domain – episodic memory – that is also highly vulnerable to aging. Given the proposed role of differentiation in age-related cognitive decline, we predicted there would be a stronger link between dedifferentiation and episodic memory performance in older than in younger adults. Young (18-30 yrs) and older (64-75 yrs) male and female humans underwent fMRI scanning while viewing images of objects and scenes prior to a subsequent recognition memory test. We computed a differentiation index in two regions-of-interest (ROIs): parahippocampal place area (PPA) and lateral occipital complex (LOC). This index quantified the selectivity of the BOLD response to an ROI’s preferred versus non-preferred category (scenes for PPA, objects for LOC). The differentiation index in the PPA, but not the LOC, was lower in older than in younger adults. Additionally, the PPA differentiation index predicted recognition memory performance for the studied items. This relationship was independent of and not moderated by age. The PPA differentiation index also predicted performance on a latent ‘fluency’ factor derived from a neuropsychological test battery; this relationship was also age invariant. These findings suggest that two independent factors, one associated with age, and the other with cognitive performance, drive neural differentiation.

**Significance Statement:** Aging is associated with neural dedifferentiation – reduced neural selectivity in ‘category selective’ cortical brain regions – which has been proposed to mediate cognitive aging. Here, we examined whether neural differentiation is predictive of episodic memory performance, and whether the relationship is moderated by age. A neural differentiation index was estimated for scene-(PPA) and object-(LOC) selective cortical regions while participants studied images for a subsequent memory test. Age related reductions were observed for the PPA, but not the LOC, differentiation index. Importantly, the PPA differentiation index demonstrated age invariant correlations with subsequent memory performance and a fluency factor derived from a neuropsychological battery. Together, these findings suggest that neural differentiation is associated with two independent factors: age and cognitive performance.

## Introduction

Healthy aging is accompanied by numerous structural (Raz et al., 2005) and functional (Spreng et al., 2010) brain changes believed to contribute to age-related cognitive decline (Raz and Rodrigue, 2006). Of relevance here is research demonstrating that increasing age is associated with reduced neural differentiation, or reduced selectivity of cortical regions sensitive to a specific class of stimuli (Park et al., 2004). A*ge-related neural dedifferentiation* has been most commonly identified in the ventral visual cortex (Grady et al., 1994; Park et al., 2004, 2010, 2012;Chee et al., 2006; Payer et al., 2006; Voss et al., 2008; Carp et al., 2011b; Kleemeyer et al., 2017; also see Berron et al., 2018), although the pattern has also been observed in auditory (Du et al., 2016) and motor cortex (Carp et al., 2011a). Neural dedifferentiation is believed to play an important role in cognitive aging (Li et al., 2001; Li and Sikström, 2002; Goh, 2011). Consistent with this proposal, measures of neural dedifferentiation have been reported to correlate negatively with cognitive performance in healthy older adults (Park et al., 2010; Du et al., 2016).

Here, we examine the proposal that neural dedifferentiation contributes to age differences in episodic memory (St-Laurent et al., 2014; Zheng et al., 2018). Healthy aging is associated with disproportionate reductions in the ability to recollect details about past events (for review, see Koen and Yonelinas, 2014; Schoemaker et al., 2014), and this deficit is largely attributed to reduced efficacy of encoding processes (Craik, 1986; Craik and Rose, 2012; Friedman and Johnson, 2014). Prior work investigating the relationship between neural dedifferentiation and memory encoding has focused on the fidelity of neural patterns across repeated instances of a given item *within* a stimulus category (St-Laurent et al., 2014; Zheng et al., 2018). The results from these studies are mixed as to whether neural dedifferentiation during encoding might contribute to age differences in memory performance. Here, we focus on indices of neural dedifferentiation measured across different stimulus categories (i.e., objects and scenes; cf.Park et al., 2004) during memory encoding, and whether these indices predict subsequent memory performance.

Participants incidentally encoded images of objects and scenes for a subsequent memory test while undergoing fMRI (see Figure 1). Objects and scenes were selected as stimuli because they selectively engage distinct cortical regions in the ventral visual cortex. Specifically, relative to scenes, viewing images of single objects engages the lateral occipital complex (LOC; Grill-Spector et al., 2001). In contrast, viewing images of scenes activates posterior parahippocampal and adjacent fusiform cortex – the ‘parahippocampal place area’ (PPA;Epstein and Kanwisher, 1998). We examined age differences in neural differentiation with a differentiation index computed from individual trial BOLD responses to objects and scenes in the LOC and PPA (Voss et al., 2008). This index reflects the scaled difference between a region-of-interest’s (ROI’s) BOLD response to a preferred (e.g., scenes in the PPA) and not preferred (e.g., objects in the PPA) stimulus category (see Materials and Methods). In a complementary analysis, neural differentiation was also examined with multi-voxel pattern analysis (cf. Carp et al., 2011). We examined the relationship between neural differentiation and two measures of memory performance, namely item recognition and source recall. Our prediction was that higher values of neural differentiation, which are indicative of increased levels of neural selectivity (Voss et al., 2008), would predict higher performance on a subsequent memory test by virtue of the mnemonic benefit associated from encoding relatively distinctive information (e.g., Murdock Jr., 1960; Lockhart et al., 1976; Hunt, 1995). Like prior research (Park et al., 2010), we also examined whether neural differentiation was associated with neuropsychological test performance. If neural dedifferentiation contributes disproportionately to memory performance (and, perhaps, performance in other cognitive domains) in older adults, differentiation should be more strongly correlated with performance in an older relative to younger participants.

## Materials and Methods

### Ethics Statement

The Institutional Review Board of the University of Texas at Dallas approved the experimental procedures described below. All participants provided written informed consent prior to participation.

### Experimental Design and Statistical Analysis

As will be elaborated in the remainder of the Materials and Methods, the main independent variables in this experiment included age group (young versus older), image type (scene versus object), and region of interest (PPA versus LOC). Results from all analyses were considered significant at *p* < .05.

Statistical analyses were conducted with *R* software (R Core Team, 2017). ANOVAs were conducted using the *afex* package (Singmann et al., 2016) and the Greenhouse-Geisser procedure (Greenhouse and Geisser, 1959) was used to correct the degrees of freedom for non-sphericity in the ANOVAs when necessary. Post-hoc tests on significant effects from the ANOVAs were conducted using the *lsmeans* package (Lenth, 2016) with degrees of freedom estimated using the Satterthwaite (1946) approximation. Effect size measures for results from the ANOVAs are reported as partial-*η*^2^ (Cohen, 1988). Linear regression models were implemented using the *lm* function in the base *R* library. Principal components analysis (PCA;Hotelling, 1933; Abdi and Williams, 2008) was conducted using the *psych* package (Revelle, 2017).

### Participants

A sample of 24 young and 24 older participants contributed to the data reported here. Participants were recruited from the University of Texas at Dallas and the greater Dallas metropolitan area and received monetary compensation ($30/hour). Table 1 reports participant demographics and neuropsychological test performance. All participants were right-handed and reported having normal or corrected-to-normal vision and no contraindications to MRI scanning. Exclusion criteria included a history of cardiovascular disease (other than treated hypertension), diabetes, psychiatric disorder, illness or trauma affecting the central nervous system, substance abuse, and self-reported current or recent use of psychotropic medication or sleeping aids. All participants scored 27 or more on the Mini-Mental State Examination (MMSE;Folstein et al., 1975) and scored within the normal range for their age group on a battery of neuropsychological tests.

**Table 1.** Demographic and neuropsychological test data for young and older adults.

|  | Young Adults | Older Adults | *p*-value |
| --- | --- | --- | --- |
| N | 24 | 24 | - |
| Age | 23.04 (3.46) | 68.92 (3.23) | - |
| Sex | 12/12 | 12/12 | - |
| Education | 15.92 (2.22) | 17.12 (2.23) | .067 |
| MMSE | 29.54 (0.59) | 29.42 (0.93) | .581 |
| CVLT Short Delay – Free | 13.08 (1.79) | 10.83 (2.84) | .002 |
| CVLT Short Delay – Cued | 13.67 (1.81) | 12.33 (2.32) | .032 |
| CVLT Long Delay – Free | 13.54 (2.06) | 10.71 (2.91) | < .001 |
| CVLT Long Delay – Cued | 14.12 (1.62) | 12.33 (2.46) | .005 |
| CVLT Recognition – Hits | 15.42 (0.83) | 15.04 (1.00) | .164 |
| CVLT Recognition – False Alarms | 0.46 (0.66) | 2.67 (2.08) | < .001 |
| Logical Memory I | 30.62 (4.95) | 26.71 (5.09) | .010 |
| Logical Memory II | 28.12 (5.78) | 23.25 (5.72) | .005 |
| Digit Span Total<sup>1</sup> | 21.04 (4.53) | 17.58 (2.41) | .002 |
| SDMT | 65.38 (13.99) | 47.21 (7.53) | < .001 |
| Trails A (secs) | 21.43 (7.97) | 30.76 (10.77) | .001 |
| Trails B (secs) | 47.54 (19.53) | 69.11 (24.64) | .002 |
| F-A-S Total | 48.29 (10.97) | 45.96 (11.65) | .479 |
| Category Fluency (Animals) | 24.58 (5.67) | 21.08 (4.82) | .026 |
| WTAR (Raw) | 41.42 (3.44) | 43.62 (4.44) | .061 |
| Raven's (List 1) | 11.08 (.97) | 9.50 (2.23) | .003 |
| Visual Acuity (logMar)<sup>2</sup> | -.11 (.10) | .06 (.11) | < .001 |
| Speed Factor (RC1)<sup>3</sup> | -.64 (.67) | .33 (.75) | < .001 |
| Memory Factor (RC2) | .55 (.73) | -.62 (1.00) | < .001 |
| Crystallized Intelligence Factor (RC3) | .00 (.79) | .08 (.93) | .751 |
| Fluency Factor (RC4) | .07 (.89) | -.21 (.72) | .257 |

*Note.* Standard deviations are reported in parentheses. The *p*-values were obtained from Welch *t*-tests comparing young and older adults. <sup>1</sup>Digit span total equals the sum of forward and backward span. <sup>2</sup>Lower logMAR scores indicate better visual acuity. <sup>3</sup>Negative factors on the speed factor (RC1) correspond to higher performance on measures of processing speed (e.g., shorter time to complete Trails A or B), whereas for other factors higher performance is indicated by higher scores. MMSE = Mini-mental State Exam; CVLT = California Verbal Learning Test II; SDMT = Symbol-Digit Modalities Test; WTAR = Wechsler Test of Adult Reading

Data from an additional 4 participants were excluded from the analyses reported here for the following reasons: 1 young adult male and 1 older adult male were excluded due to excessive in-scanner motion (> 8 mm frame displacement) and 2 older adult males were excluded for providing 2 or fewer source correct trials (see below).

Many participants in the present study participated in prior studies reported by our laboratory. Specifically, 18 young (10 females) and 16 older (4 females) participated in an ERP study reported by Koen and colleagues (2018). Additionally, 2 older adults (1 female) participated in a prior fMRI experiment reported by de Chastelaine and colleagues (2016).

### Neuropsychological Test Battery

Participants completed a neuropsychological test battery on a separate day prior to the fMRI study. The battery included the MMSE, California Verbal Learning Test-II (CVLT;Delis et al., 2000), the symbol digit modalities test (Smith, 1982), forward and backward digit span subtests of the Wechsler Adult Intelligence Scale – Revised (Wechsler, 1981), trail making tests A and B (Reitan and Wolfson, 1985), the F-A-S subtest of the Neurosensory Center Comprehensive Evaluation for Aphasia (Spreen and Benton, 1977), the category fluency test for animals (Benton, 1968), Wechsler test of adult reading (WTAR; Wechsler, 2001), the logical memory subtest of the Wechsler Memory Scale (Wechsler, 2009), and List 1 of the Raven’s Progressive Matrices (Raven et al., 2000). Volunteers were excluded from participating in the fMRI study if (1) one or more of the memory measures (i.e., CVLT or logical memory) were more than 1.5 standard deviations below the age- and education-adjusted mean, (2) they had a standard score below 100 on the WTAR, or (3) two or more scores on non-memory tests were 1.5 standard deviations below the mean (see below for the dependent measures that were used).

### Neuropsychological Data Analysis

The scores on the neuropsychological test battery were reduced to factor scores based on PCA applied to a prior dataset from our laboratory that included young, middle, and older adults (de Chastelaine et al., 2016). Principal components with eigenvalues > 1 were kept and rotated using Varimax rotation (Kaiser, 1958). The following variables were included in the PCA model: CVLT composite recall measure (i.e., average number of words recalled on the short- and long-delay free- and cued-recall tests), number of CVLT recognition hits, number of CVLT recognition false alarms, a logical memory composite recall measure (i.e., average of immediate and delayed recalls), completion time for both trails A and B, number of valid responses on the SDMT, F-A-S, and Raven’s, and estimated full-scale intelligence quotient derived from the WTAR. The first four components were retained and explained 64.1% of the variance in the data prior to rotation. The rotated components (RC) broadly correspond to factors representing processing speed (RC1), memory (RC2), crystallized intelligence (RC3), and fluency (RC4). The weights for the rotated factors from this prior data set are shown in Table 2. These weights were applied to the identical variables in the present data set to extract factor scores for the analyses reported here.

**Table 2.**
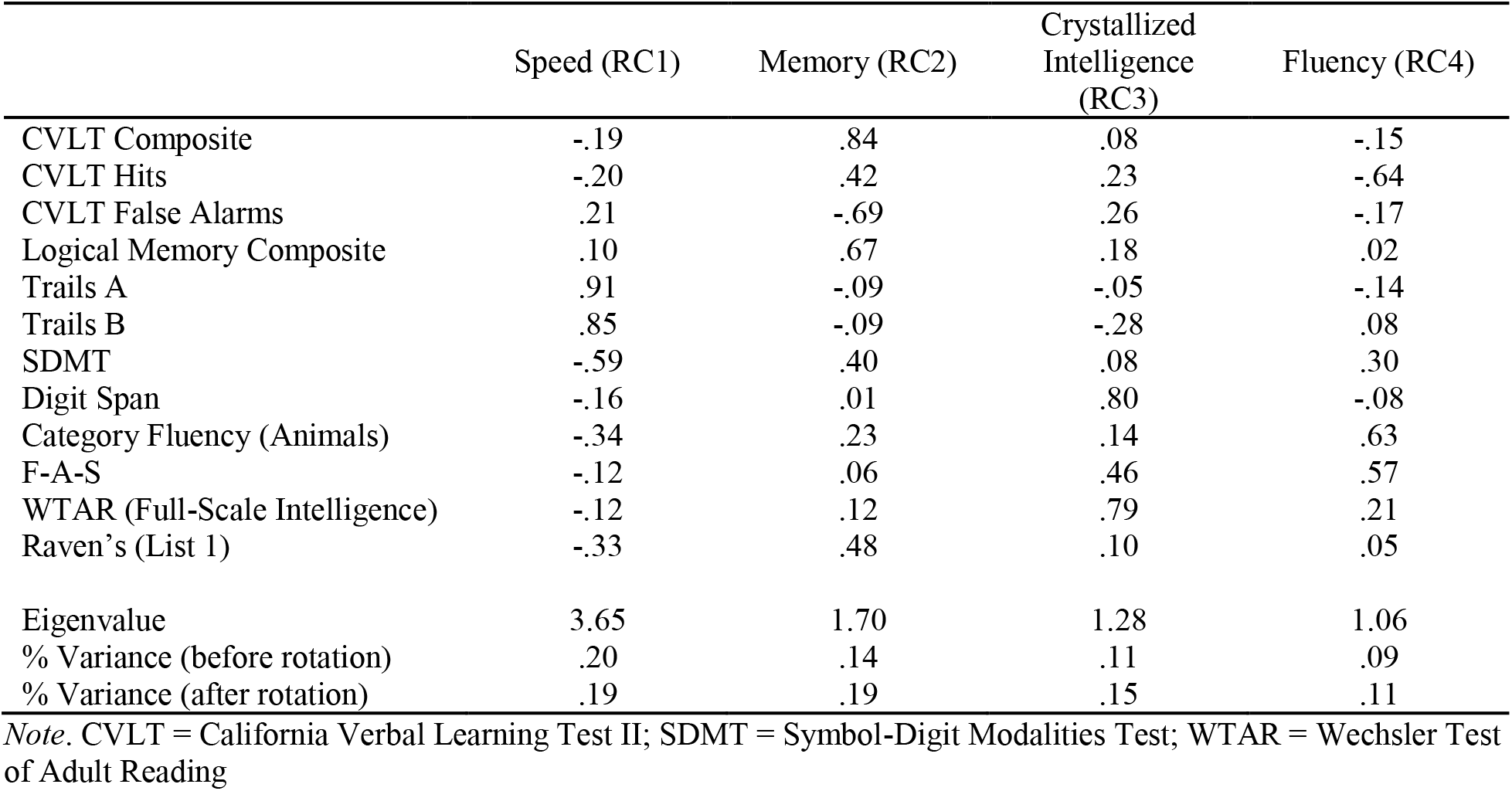
Rotated factor loadings from the PCA (with Varimax rotation) of the neuropsychological test data reported by de Chastelaine et al. (2016).

|  | Speed (RC1) | Memory (RC2) | Crystallized<br>Intelligence<br>(RC3) | Fluency (RC4) |
| --- | --- | --- | --- | --- |
| CVLT Composite | -.19 | .84 | .08 | -.15 |
| CVLT Hits | -.20 | .42 | .23 | -.64 |
| CVLT False Alarms | .21 | -.69 | .26 | -.17 |
| Logical Memory Composite | .10 | .67 | .18 | .02 |
| Trails A | .91 | -.09 | -.05 | -.14 |
| Trails B | .85 | -.09 | -.28 | .08 |
| SDMT | -.59 | .40 | .08 | .30 |
| Digit Span | -.16 | .01 | .80 | -.08 |
| Category Fluency (Animals) | -.34 | .23 | .14 | .63 |
| F-A-S | -.12 | .06 | .46 | .57 |
| WTAR (Full-Scale Intelligence) | -.12 | .12 | .79 | .21 |
| Raven's (List 1) | -.33 | .48 | .10 | .05 |
| Eigenvalue | 3.65 | 1.70 | 1.28 | 1.06 |
| % Variance (before rotation) | .20 | .14 | .11 | .09 |
| % Variance (after rotation) | .19 | .19 | .15 | .11 |

*Note.* CVLT = California Verbal Learning Test II; SDMT = Symbol-Digit Modalities Test; WTAR = Wechsler Test of Adult Reading

### Visual Acuity Assessment

Participants completed a visual acuity test using ETDRS charts (Precision Vision, La Salle, Illinois) during the neuropsychological test session. Visual acuity was measured separately for the left and right eyes, as well as with both eyes using the logMAR metric (Ferris et al., 1982; Bailey and Lovie-Kitchin, 2013). A different eye chart was used for each of the three tests. Participants prescribed corrective lenses wore them during the visual acuity test. Note that only the results from the visual acuity measured with both eyes is reported (see Table 1).

### Materials and Apparatus

Stimuli were presented using Cogent 2000 software (www.vislab.ucl.ac.uk/cogent_2000.php) implemented in Matlab 2011b (www.mathworks.com). Stimuli in the scanned study phases were projected to a screen mounted at the rear of the magnet bore and viewed through a mirror mounted on the head coil. Responses during the study sessions were entered using two four-button MRI compatible response boxes (one for each hand). The test phase was completed on a laptop computer outside the scanner. The monitor resolution setting for both the study and test phases was set at 1024 x 768 pixels. All stimuli were presented on a grey background (RGB values of 102, 101 and,99).

The critical stimuli comprised 360 images obtained from a variety of internet sources. Half of the images were pictures of scenes and the remaining half were pictures of common objects. The 180 scenes comprised 90 rural (i.e., natural) scenes and 90 urban (i.e., manmade) scenes. The scenes contained objects (e.g., trees, cars, buildings, etc.), and we attempted to minimize overlap between the objects depicted in the scenes and the object images. The scenes were scaled and cropped to 256 x 256 pixels.

The 180 objects comprised 90 images of natural objects (e.g., food items, animals, plants) and 90 images of manmade objects (e.g., tools, vehicles, furniture). The object images were overlaid and centered on a light grey background (RGB values of 175, 180, and 184) with dimensions of 256 x 256 pixels. Note that the background color for the object images differed from the background of the monitor. The purpose of this was to roughly equate the area of the monitor subtended by the object and scene images.

The above-described images were used to create 24 stimulus sets that were yoked across young and older participants. Each stimulus set comprised a random selection of 120 objects and 120 scenes that served as study items. The 120 images of each type were divided into 5 groups of 24, and each group was randomly assigned to one of the five scanned study phases. Half of the objects and scenes in each study session were assigned to each of the two different possible judgments in the study phase (Pleasantness and Movie; see below). The test stimuli comprised all the images from the study phase along with the remaining 60 objects and 60 scenes, which served as new items. All stimulus lists were pseudorandomized such that there were no more than three consecutive presentations of objects or scenes and no more than three consecutive Pleasantness or Movie judgments.

An additional 16 objects and 16 scenes with similar characteristics to those described above served as practice stimuli. The images in each practice list were the same for all participants. There were 3 practice study lists (self-paced, speeded, real; see below), each comprising 8 images (4 objects, 4 scenes). A practice test list was also created and comprised the images from the speeded and real practice study phases (old items) and 8 images (4 objects, 4 scenes) as new items.

### Procedure

#### Overview

The experiment was completed across two sessions on different days, with the neuropsychological test battery completed in the first session, and the experimental fMRI session completed in the second session. In the fMRI session, participants first completed a face-viewing task in which they pressed a button when an inverted face appeared among a sequence of upright faces. The face-viewing task is not discussed further here and will be the subject of a separate report. Following the face-viewing task, participants completed the study phase of the experiment described here, followed by a test phase administered outside of the scanner (see Figure 1).

**Figure 1.**
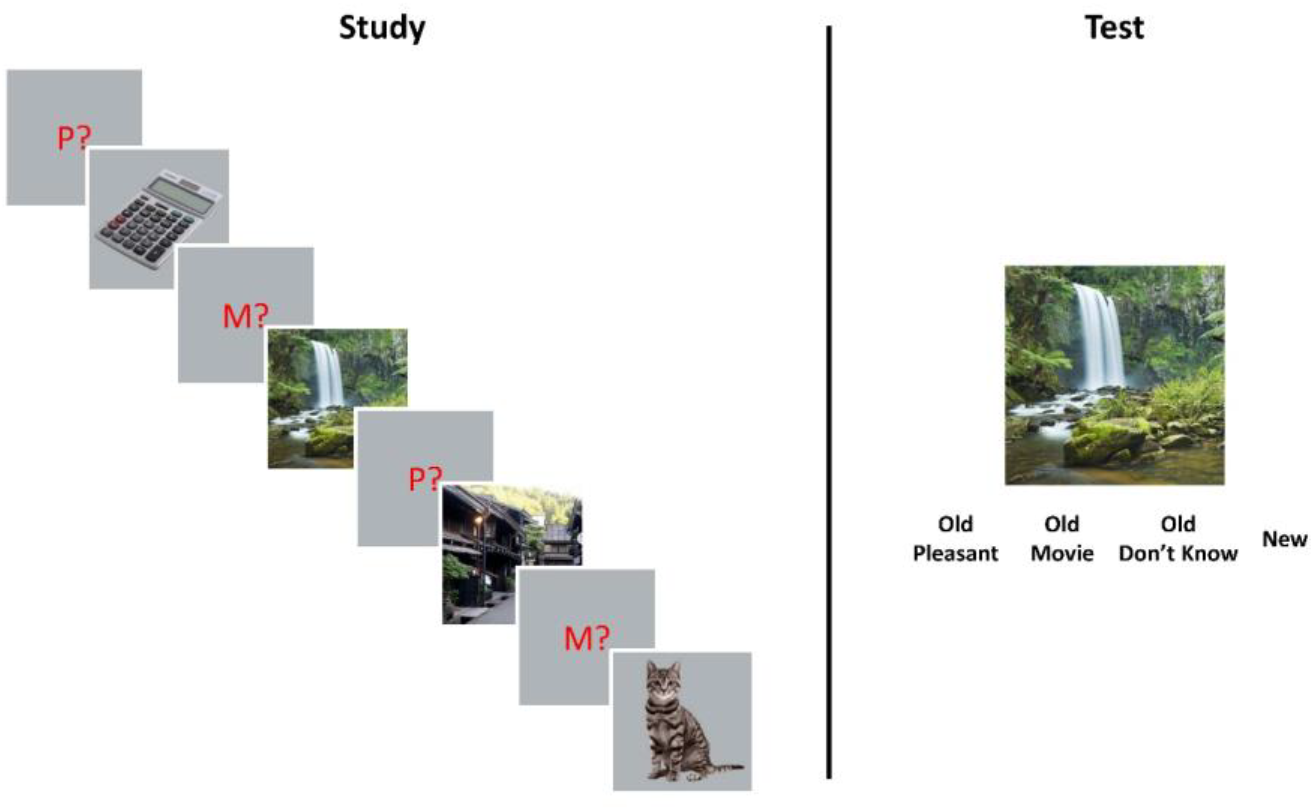
Schematic overview of the memory task. Participants studied an intermixed list of object and scene images under intentional encoding instructions while undergoing fMRI scanning. Each image was preceded by a task cue that instructed participants to rate the image for pleasantness (P?) or to determine which movie genre the image was best associated with (movie, M?). There were a total of 5 scanned study phases. After the final study phase, an out-of-scanner recognition memory test was administered. The test phase comprised the studied objects and scenes intermixed with new images. Participants were instructed to select one of four memory judgments for each image. The four judgments comprised options for whether participants had high confidence both that they studied the image and could recollect the study task (Old Pleasant and Old Movie responses), had high confidence that they studied the image but were had low confidence in their memory for or could not remember the study task (Old Don’t Know response), or if they did not have high confidence that the image was studied (New response). Two measures of memory performance were obtained from the test phase: item recognition and recall of the encoding task (i.e., source recall).

#### Study Phase

Participants completed the study phase during five consecutive fMRI scanning sessions. The study phase was completed under intentional encoding conditions with specific reference to the nature of the subsequent memory test.

The sequence and timing for each trial was as follows: get ready signal (green fixation cross for 500 ms), task cue (red ‘P?’ or ‘M?’ for 500 ms), study image (object or scene for 2000 ms), and white fixation (1750 ms). The task cue informed participants which one of two judgments they should make about the following image. Images preceded by a ‘P?’ (Pleasantness) required participants to rate how pleasant they found the image using the following scale: ‘Very’, ‘Moderate’, or ‘Not at all’. Images preceded by a ‘M?’ (Movie) required participants to determine which movie genre they believed was best associated with the object or scene. There were three options for this judgment: ‘Action’, ‘Horror’, or ‘Comedy’. The response options for the cued judgment always appeared below the image.

Participants were instructed to enter their responses quickly, and to attempt to do so while the image was on the screen. Responses were entered with the index, middle and ring fingers (respectively for the order of response options listed previously), and were accepted until the beginning of the next trial. Responses for one judgment were entered with the right hand and responses for the other judgment were entered with the left hand. The hand assigned to each question was counterbalanced across participants. The instructions emphasized that responding with the incorrect hand for a cued judgment counted as an incorrect response.

In addition to the critical trials, there were 24 null trials dispersed throughout each of the 5 scanned study sessions. The null trials displayed a white fixation cross for the duration of a normal trial (4750 ms) and were distributed such that 12 objects and 12 scenes were each followed by a single null trial. This was done to minimize any bias between the two image types in estimating single trial BOLD responses. Null trials never occurred consecutively, resulting in stimulus onset asynchronies of either 4750 or 9500 ms for both classes of image.

#### Test Phase

The test phase commenced outside of the scanner approximately 15 minutes after the completion of the final study phase. Participants were shown images one at a time and required to judge if the image was presented in the study phase while they were in the scanner and, if so, which of the two encoding judgments they had made when they initially encountered the image. These two mnemonic decisions were combined into a single judgment with four possible options: ‘Old-Pleasant’, ‘Old-Movie’, ‘Old-Don’t Know’, ‘New’. A ‘New’ response was required if the image was believed to be new or if participants had a low level of confidence that the image was from the study list. An ‘Old-Pleasant’ or ‘Old-Movie’ response required participants to have high confidence that they studied the image and high confidence in their memory for the judgment made when the image was studied. Participants were instructed to respond ‘Old-Don’t Know’ if they had high confidence they studied the image but had low confidence in or were unable to remember the encoding judgment.

Responses were entered on the keyboard by pressing the ‘d’, ‘f’, ‘j’, and ‘k’ key, and these keys were labeled ‘Old-Pleasant’, ‘Old-Movie’, ‘Old-Don’t Know’, and ‘New’, respectively. Responses were self-paced, but participants were instructed to enter their responses quickly without sacrificing accuracy. There was a brief 500 ms white fixation cross between test trials. A short break was afforded to participants every 60 trials (totaling 5 breaks).

#### Practice Phases

Prior to MRI scanning, participants practiced both the study and test phases outside of the scanner. Practice comprised 3 study phases and a single test phase. In the self-paced practice phase, participants were presented with the trial sequence as described above, with the exception that the image remained on the screen until a response was entered. Following a response, participants received feedback as to whether they responded to the correct judgment (i.e., whether they entered their judgment using the assigned hand for the Pleasantness or Movie judgments). The trial was repeated in the event the incorrect hand was used, and this occurred until the correct hand was used. The aim of this self-paced practice phase was to familiarize participants with responding to each type of judgment using the correct hand.

Next, participants completed a speeded practice phase. This phase was identical to the self-paced practice described above, with the exception that the image remained on the screen only for 2000 ms. Participants were required to enter their response within this time window, otherwise they were given feedback that they did not enter a response in the allotted time. As with the self-paced practice study phase, a trial was repeated until the correct hand was used and a response was entered in the allotted time. The aim of this second practice study phase was to reinforce responding with the correct hand and to give participants experience with responding quickly. No null trials were included in the self-paced and speeded practice study phases. The final ‘real’ practice study phase mirrored the procedure for the study phase proper described above and included 4 null trials.

After the final practice study phase, participants completed the practice test phase. This mirrored the procedure for the test phase proper with the exception that no breaks were provided.

### Behavioral Data Analysis

Trials that received no response or a response with the incorrect hand during the study phase were excluded from the analysis. Both study and test trials were binned according to the four possible test response outcomes: item hit with a correct source judgment, item hit with an incorrect source judgment, item hit accompanied by a don’t know response for the source judgment, and item misses. Note that new items do not have a source correct judgment, thus false alarms (i.e., incorrect ‘old’ responses to new images) were only classified as source incorrect or source don’t know trials. The three behavioral dependent measures analyzed included study reaction time (RT), item recognition accuracy, and source memory accuracy. Study RT was computed for each participant as the median RT for each image type and subsequent memory combination. There were three subsequent memory bins: source correct (SC), source incorrect/don’t know (SIDK), and item misses (Miss). Study RT was analyzed with a 2 (Age Group: Young, Older) X 2 (Image Type: Object, Scene) X 3 (Subsequent Memory: SC, SIDK, Miss) mixed-factorial ANOVA.

Item recognition accuracy was computed as the difference between the hit rate to studied images (regardless of source memory accuracy) and the false alarm rate to new images. Source memory was computed using a single-high threshold model (Snodgrass and Corwin, 1988) that accounts for the ‘guess rate’ (e.g., Mattson et al., 2014). Source accuracy was computed as follows:

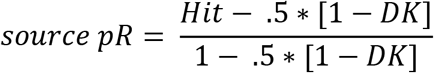

The *Hit* and *DK* variables in the above formula refer to the proportion of correct ‘old’ responses (i.e., hits) accompanied by an accurate or don’t know source memory judgments, respectively. The item and source memory scores were submitted to separate 2 (Age Group: Young, Older) X 2 (Image Type: Object, Scene) mixed-factorial ANOVA.

### Identification of PPA and LOC Regions-of-Interest

The analyses of the fMRI data focused on two regions-of-interest (ROIs) that show selective responses to scenes and objects, respectively: the parahippocampal place area (PPA;Epstein and Kanwisher, 1998) and lateral occipital complex (LOC;Grill-Spector et al., 2001). We identified these ROIs bilaterally using unpublished data from our laboratory obtained from a sample of 22 participants (14 young and 8 older adults) who volunteered for a previous study (see Figure 2A). Note that 1 young and 2 older participants from this unpublished study overlapped with the participants reported here. The 22 participants viewed images of faces, scenes, and articles of clothing (objects) in a mini-block design (e.g., Johnson et al., 2009; McDuff et al., 2009; Wang et al., 2016) while providing a pleasantness rating for each image. PPA and LOC ROIs were obtained from a second-level general linear model (GLM) contrasting the BOLD response between scenes and objects. The two one-sided contrasts were thresholded at a family-wise error (FWE) corrected threshold of *p* < .05, and were inclusively masked using anatomical labels from the Neuroinformatics atlas included with SPM12. The bilateral PPA ROI comprised 223 voxels (108 voxels in the left hemisphere) identified by the scene > object contrast anatomically masked with the bilateral parahippocampal and fusiform gyri. The bilateral LOC ROI comprised 225 voxels (98 voxels in the left hemisphere) identified by the object > scene contrast anatomically masked inferior and middle occipital gyrus ROIs defined by the Neuroinformatics atlas. The PPA and LOC ROIs used for the present study are depicted in Figure 2A. Additionally, Figure 2B shows the statistical maps from the scene > object (warm colors) and object > scene (cool colors) contrasts from a 2^nd^ level GLM of our unpublished data set without the anatomical inclusive mask. Figure 2C shows the same statistical contrast (at an identical threshold to Figure 2B) for the 24 young and 24 older adults reported here. This is included simply for comparison purposes. Note that differences in the magnitude and extent of the contrasts in Figures 2B and 2C are likely attributable to the larger sample size in the present study.

**Figure 2.**
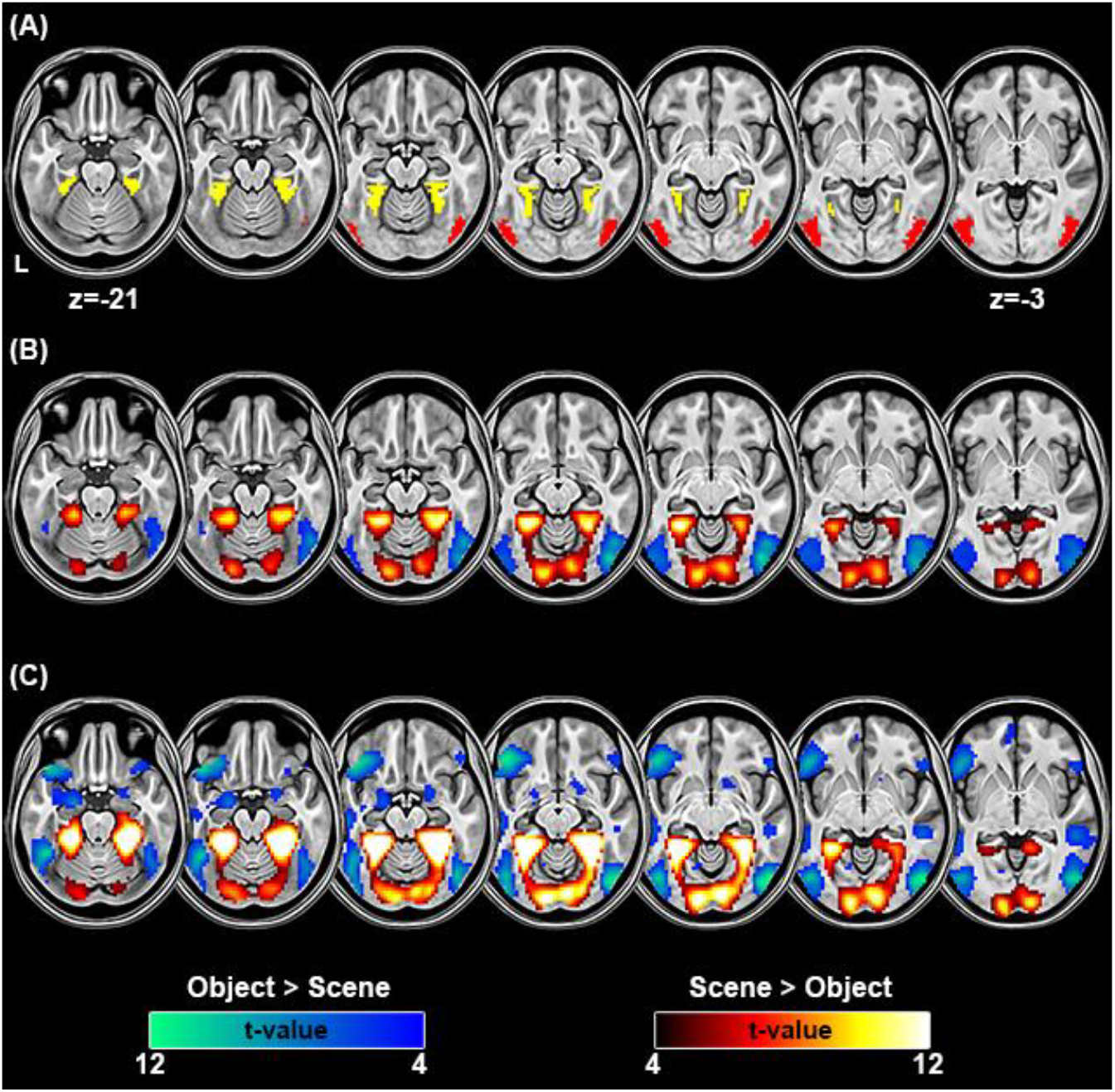
(A) Voxels comprising the regions-of-interest (ROIs) in the parahippocampal place area (PPA; yellow voxels) and lateral occipital cortex (LOC; red voxels) derived from an unpublished data set. Note that the ROIs were anatomically masked using the Neuroinformatics atlas included in SPM12. The anatomical labels for this mask included bilateral parahippocampal, fusiform, middle occipital, and inferior occipital gyri. (B) Statistical parameteric maps (SPMs) from the unpublished experiment showing the one-tailed contrasts of Scene > Objects and Objects > Scenes. (C) SPMs for the Scene > Objects and Objects > Scene contrast in the 24 young and 24 older adults in the present data (collapsed across age group). The SPMs are thresholded at FWE of *p* < .05 (FWE).

### MRI Data Acquisition

MRI data were acquired with a 3T Philips Achieva MRI scanner (Philips Medical Systems, Andover, MA, USA) equipped with a 32-channel receiver head coil. Functional images were acquired with a blood oxygenation level dependent (BOLD), T2*-weighted echoplanar imaging (EPI) sequence (SENSE factor = 1.5, flip angle = 70°, 80 × 80 matrix, FOV = 240 mm x 240 mm, TR = 2000 ms, TE = 30 ms, 34 ascending slices, slice thickness = 3 mm, slice gap = 1 mm), and were oriented parallel to AC-PC. Five “dummy” scans were acquired at the start of each EPI session and discarded to allow for equilibration of tissue magnetization. A total of 180 functional volumes were acquired during each study session, for a total of 900 brain volumes. T1-weighted images (MPRAGE sequence, 240 × 240 matrix, 1 mm isotropic voxels) were acquired for anatomical reference following prior to the first study session.

### Formation of Study Specific MNI Templates

A sample specific EPI template was created using the mean EPI image from all participants included in the analysis following previously published procedures (de Chastelaine et al., 2011, 2016). Each participant’s mean EPI image was first normalized to the standard EPI template in SPM12, and the spatially normalized images were then averaged within age group to create a young and older adult EPI template. The final template was created by averaging the two age-specific templates.

### fMRI Preprocessing

The functional data were preprocessed with Statistical Parametric Mapping (SPM12, Wellcome Department of Cognitive Neurology, London, UK) implemented in Matlab 2017b (The Mathworks, Inc., USA). The functional data were reoriented, subjected to a two-pass realignment procedure whereby images were first realigned to the first image of a session and then realigned to a mean EPI image, and corrected for slice acquisition time differences using sinc interpolation with reference to the middle slice. Finally, images were spatially normalized to a study specific EPI template (see Creation of Study Specific MNI Templates below), and smoothed with an 8mm full-width at half-maximum kernel.

The data from the five study sessions were concatenated and subjected to a least-squares-all (LSA) GLM to estimate the BOLD response to individual trials (Rissman et al., 2004; Mumford et al., 2014). Events were modeled as a 2 s-duration boxcar convolved with a canonical HRF. Covariates of no interest in this first level model included the 6 rigid body motion parameters estimated from the realignment procedure and 4 session specific means (for sessions 2-5).

### Differentiation Index Analysis

We computed a differentiation index for the PPA and LOC ROIs (see Identifying PPA and LOC Regions-of-Interest). For each trial, we extracted the average BOLD amplitude separately for each ROI (collapsed across hemisphere). These individual trial values were used to compute separate differentiation indices for each bilateral ROI using a similar formula to that employed by Voss and colleagues (2008). The index is essentially a discrimination metric similar to the *d’* signal detection measure (Macmillan and Creelman, 2005), and was computed using the following formula:

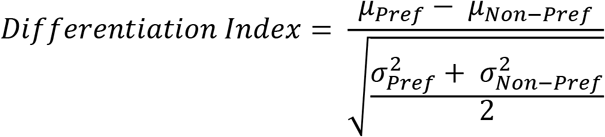

In the above equation, *μ_pref_* and 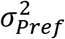 refer to the across-trial mean and variance, respectively, of the BOLD response to an ROI’s preferred image type. The *μ_Non−pref_* and 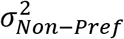 terms refer to the across-trial mean and variance, respectively, of the non-preferred image type. For the PPA, scenes were designated as the preferred image type and objects as the non-preferred image type, and this designation was reversed for the LOC.

Positive values of the differentiation index reflect higher ‘selectivity’ of responding to an ROI’s preferred image type. We note two aspects of this index that bear mention. First, and importantly, the differentiation index is insensitive to across-participant variability in the hemodynamic response function and, therefore, is unbiased by putative systematic age-differences in such factors as cerebral vascular reactivity (see, for example,Liu et al., 2013). Second, the index is a metric of category selectivity, and does not measure selectivity at the ‘item level’ (for potential approaches to item level distinctiveness, see Goh et al., 2010; St-Laurent et al., 2014; Xue et al., 2010). The differentiation index data were subjected to a 2 (Age Group) X 2 (ROI: PPA, LOC) mixed factorial ANOVA.

An additional ANOVA of the differentiation index data was conducted in which subsequent memory bin (SC, SIDK, Miss) was included as a factor. This ANOVA produced identical results to the 2 × 2 ANOVA described above, with no effects involving subsequent memory. Thus, for simplicity’s sake, we focus below on the differentiation index computed across all trials regardless of subsequent memory judgment.

The differentiation index is ambiguous with respect to whether a group difference, if any, is driven by reduced BOLD signal for the preferred image type (i.e., neural attenuation), an increase in BOLD signal for the non-preferred image type (i.e., neural broadening), or by both effects (cf.Park et al., 2012). To investigate this issue, we also examined the mean BOLD responses elicited by each image type within the two ROIs using a 2 (Age Group) X 2 (ROI) X 2 (Image Type: Object, Scene) mixed factorial ANOVA.

A primary goal of the present study was to examine whether neural differentiation during encoding is predictive of subsequent memory performance. We addressed this issue by computing across-participant correlations between the PPA and LOC differentiation indices, and performance on the experimental memory task (i.e., item recognition and source memory scores). Additionally, we computed partial correlations between these indices after controlling for several relevant variables, including age group, item or source memory performance (when source and item memory were in the zero-order correlation, respectively), and visual acuity.

For clarity, we focus here on the partial correlations. Results from multiple regression analyses led to conclusions identical to those derived from the partial correlation analyses reported below. Of importance, the inclusion of an interaction term between age and the neural differentiation indices in the regression models did not significantly increase the amount of explained variance compared to models with only age group and differentiation indices as predictors, *F*’s(1,44) < 2.83, *p* ≥ .100, nor did the regression coefficients for the interaction terms approach significance. Thus, we found no support that any of the reported correlations between differentiation indices and memory performance were moderated by age group. Moreover, in the analyses reported below, partial correlations were computed after averaging the memory measures across image type, as there was no indication that the effects of interest were moderated by this variable. Specifically, in a multilevel regression conducted with the *lmerTest* package in *R* (Kuznetsova et al., 2015), no interaction term that involved that variable of image type approached significance, all regression coefficients *p*’s ≥ .136. The full results from these multiple and multilevel regression analyses are available from the first author upon request.

In addition to the correlation analyses involving memory performance, we also examined the relationship between the differentiation indices and the extracted factor scores for the neuropsychological test battery (see Analysis of Neuropsychological Data), again with partial correlations. Importantly, as with the two memory measures, multiple regression provided no evidence that the relationship between any of the factor scores and the differentiation indices were moderated by age group, *F*’s(1,44) < 1.66, *p* ≥ .204. A multilevel regression model including a factor for the four RC scores led to identical conclusions to those derived from the partial correlations reported below. These regression analyses also are available from the first author upon request.

### Pattern Similarity Analysis

To complement the analyses of the univariate differentiation index described above, we also conducted a pattern similarity analysis (PSA;Kriegeskorte et al., 2008). All similarity computations were conducted on single-trial beta weights (see above) and were based on Fisher-*z* transformed Pearson’s correlation coefficients. A within-minus-between (henceforth within-between) similarity metric was computed separately for each ROI with the preferred and non-preferred image category serving as the within and between measure, respectively. For the PPA, the within-category measure was the average across-voxel similarity between a given scene trial with all other scene trials. The between-category similarity measure was the average correlation between a given scene trial and all object trials. For each scene trial in the PPA, the within-between measure was computed as the difference between the above described within and between similarity metrics. A summary measure for a participant was computed by averaging all of the trial-wise within-between measures. The same approach was used to compute the within-between similarity metric for the LOC, except that object trials were used for the within-category measures, and scene trials provided the between-category measures. We refer to the metric as the ‘similarity index’. Analogous to the differentiation index described above, the similarity index is a measure of similarity at the category and not the item level. The similarity indices were subjected to a 2 (Age Group) X 2 (ROI: PPA, LOC) mixed factorial ANOVA.

As for the univariate differentiation index describe above, ANOVA of the similarity metrics that included a subsequent memory factor (SC, SIDK, Miss) revealed no effects involving subsequent memory. Therefore, we report the similarity findings collapsed across subsequent memory judgment. Further echoing the analyses of the differentiation index, we examined the associations between the pattern similarity index and memory and neuropsychological test performance and report the findings in terms of partial correlations.

Analysis using multiple regression led to identical conclusions; crucially, there was no indication that adding a term for the interaction between age group and the similarity index improved model fit beyond that obtained with models without this term, *F*’s(1,44) < 1.35, *p* ≥ .144, and nor did the regression coefficients for any of the interaction terms approach significance. Thus, we found no evidence that the correlations reported between the pattern similarity index and cognitive performance were moderated by age group.

## Results

### Neuropsychological Test Performance

The results from the different measures of the neuropsychological test battery are reported in Table 1. The pattern of age differences is essentially identical to our prior report (Koen et al., 2018), which is not surprising given the high degree of overlap between the samples (see Participants section of the Methods). There were significant effects of age, with older adults performing worse on tests assessing declarative memory, reasoning ability, category fluency, and processing speed. However, older adults were equally proficient at word reading and verbal fluency relative to young adults. Finally, as expected (e.g., Baltes and Lindenberger, 1997), older participants had lower visual acuity than younger adults.

The bottom portion of Table 1 shows extracted factor scores derived from the test (see Table 2 for the rotated PCA loadings and the Neuropsychogical Test Analysis section). Not surprisingly, and consistent with the analysis of the individual tests, there were age differences in the speed (RC1) and memory (RC2) factors. No age differences were observed for the factors corresponding to crystallized intelligence (RC3) and fluency (RC4).

### Study Reaction Time

Table 2 shows the descriptive statistics of the median RTs for the study judgments. A 2 (Age Group) X 2 (Image Type) X 3 (Subsequent Memory) mixed ANOVA revealed a main effect of subsequent memory, *F*(1.96,90.31) = 24.43, *MS*_e_ = 8705, *p* < 10^−8^, partial-*η*^2^ = .35, that was driven by faster RTs for subsequent source correct trials (*M* = 1321) relative to both source incorrect (*M* = 1399), *t*(92) = 5.86, *SE* = 13.34, *p* < 10^−4^ and item miss trials (*M* = 1404), *t*(92) = 6.23, *SE* = 13.34, *p* < 10^−4^. There was no significant difference between study RTs associated with subsequent incorrect source memory and item misses, *t*(92) = .37, *SE* = 13.34, *p* = .712. Nor were there any significant effects involving age group (all *p*’s involving Age Group > .133).

**Table 3.**
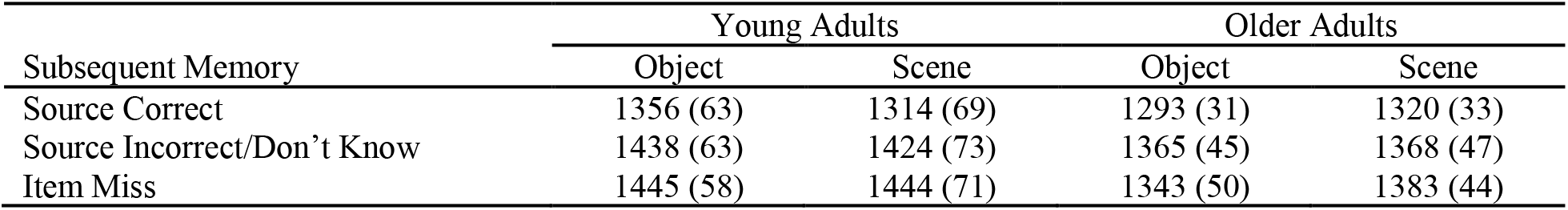
Mean (and standard errors) for the median RT (in ms) to judgments made during the study phase.

| Subsequent Memory | Young Adults |  | Older Adults |  |
| --- | --- | --- | --- | --- |
|  | Object | Scene | Object | Scene |
| Source Correct | 1356 (63) | 1314 (69) | 1293 (31) | 1320 (33) |
| Source Incorrect/Don't Know | 1438 (63) | 1424 (73) | 1365 (45) | 1368 (47) |
| Item Miss | 1445 (58) | 1444 (71) | 1343 (50) | 1383 (44) |

### Memory Performance

Table 4 shows the mean proportion of responses given to test items as a function of age group, image type, and study status (old or new), while Table 5 reports the item and source memory scores for objects and scenes in young and older adults. A 2 (Age Group) X 2 (Image Type) mixed factorial ANOVA on the item recognition measure revealed a significant main effect of image type, *F*(1,46) = 187.97, *MS*_e_ = .01, *p* < 10^−15^, partial-*η*^2^ = .80, reflecting better item recognition for objects than scenes. Although older adults (*M* = .57, *SE* = .03) demonstrated numerically lower item recognition scores than young adults (*M* = .65, *SE* = .03), the main effect of age group was not significant according to our *a priori* statistical threshold, *F*(1,46) = 3.89, *MS*_e_ = .04, *p* = .055, partial-*η*^2^ = .08. The interaction between age and image type was not significant, *F*(1,46) = 1.04, *MS*_e_ = .01, *p* = .312, partial-*η*^2^ = .02.

**Table 4.** Means (with standard errors) for the proportion of trials in each cell formed by age group, image type, and item type (old versus new) for the four possible memory response bins.

| Test Response | Young Adults |  |  |  | Older Adults |  |  |  |
| --- | --- | --- | --- | --- | --- | --- | --- | --- |
|  | Objects |  | Scenes |  | Objects |  | Scenes |  |
|  | Old | New | Old | New | Old | New | Old | New |
| Old+SC | .58 (.05) | – | .32 (.03) | – | .56 (.04) | – | .34 (.03) | – |
| Old+SI | .04 (.01) | .01 (.01) | .05 (.01) | .03 (.01) | .13 (.02) | .08 (.02) | .12 (.02) | .14 (.03) |
| Old+DK | .21 (.03) | .04 (.02) | .29 (.02) | .11 (.02) | .14 (.03) | .03 (.04) | .24 (.03) | .15 (.03) |
| New | .17 (.03) | .95 (.02) | .33 (.04) | .86 (.03) | .17 (.02) | .89 (.02) | .30 (.03) | .72 (.04) |

*Note.* It is impossible to have a source correct (SC) response for new trials. Thus, incorrect old responses to new items are classified as a source incorrect (SI) trial if participants selected one of the two encoding tasks or as a source don't know (DK) trial if participants selected the don't know response option.

An analogous 2 × 2 mixed factorial ANOVA on the source memory measure also produced a significant main effect of image type, *F*(1,46) = 105.05, *MS*_e_ = .01, *p* < 10^−12^, partial-*η*^2^ = .70 which was driven by better source memory for objects than for scenes (see Table 5). There was no significant difference in source memory accuracy between young and older adults, *F*(1,46) = .81, *MS*_e_ = .06, *p* = .372, partial-*η*^2^ = .02, and nor was there a significant interaction between age and image type, *F*(1,46) = .97, *MS*_e_ = .01, *p* = .329, partial-*η*^2^ = .02.

**Table 5.** Means (with standard errors) estimates of item and source memory discrimination.

| Age Group | Item Recognition |  | Source Memory |  |
| --- | --- | --- | --- | --- |
|  | Object | Scene | Object | Scene |
| Young Adults | .78 (.04) | .52 (.04) | .51 (.05) | .27 (.03) |
| Older Adults | .72 (.03) | .42 (.03) | .44 (.04) | .25 (.03) |

*Note.* Item recognition reflects the difference between the hit and false alarm rate regardless of source memory accuracy. Source memory was computed with the pR formula (see Behavioral Data Analysis) only for studied images attracting an accurate 'old' response.

### Differentiation Index

The results from the fMRI differentiation index are presented in Figure 3A. A 2 (Age Group) X 2 (ROI) mixed factorial ANOVA on these data produced a significant interaction, *F*(1,46) = 20.31, *MS*_e_ = .06, *p* < 10^−4^, partial-*η*^2^ = .31. The interaction was driven by significantly lower differentiation indices from the PPA in older relative to younger adults, *t*(91.71) = 5.76, *p* < 10^−4^. No age differences were observed in the LOC differentiation index, *t*(91.71) = .60, *p* = .551.

**Figure 3.**
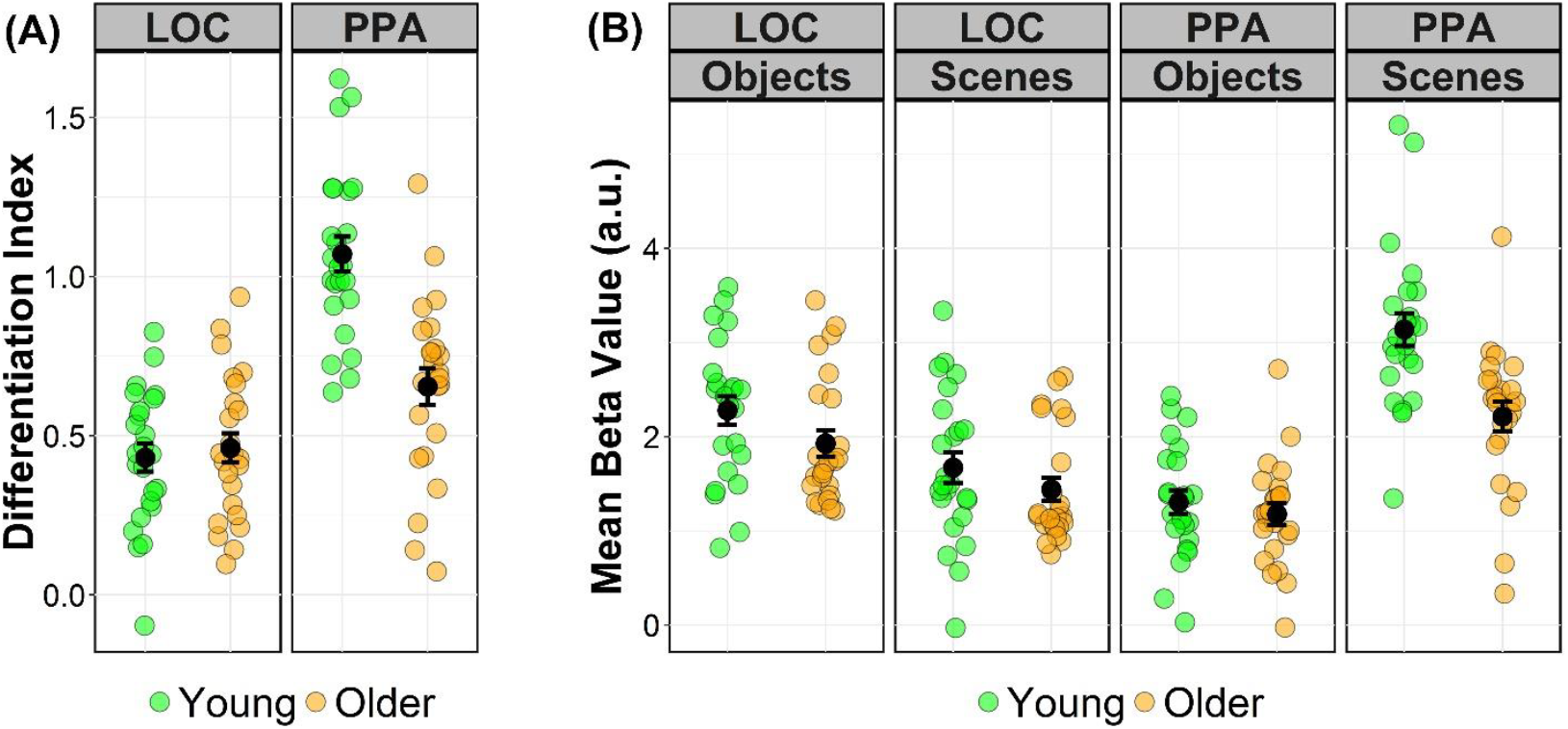
(A) Plot of the differentiation index computed from the LOC and PPA for young and older adults. (B) Plot of the across-trial mean beta-values for each image type and region of interest. Each green and orange circles represent an individual participant’s data, and the black circle represents the group mean with error bars denoting ±1 standard error of the mean.

To investigate if the age-related reduction in the PPA differentiation index resulted from reduced BOLD signal for the region’s preferred stimulus type (i.e., neural attenuation), increased BOLD signal for an ROIs non-preferred stimulus type (i.e., neural broadening), or a mixture of the two, we conducted a 2 (Age Group) X 2 (ROI) X 2 (Image Type) mixed factorial ANOVA on the mean BOLD responses (see Figure 3B). The ANOVA produced a significant three-way interaction, *F*(1,46) = 37.76, *MS*_e_ = .45, *p* < 10^−6^, partial-*η*^2^ = .31. Post-hoc tests demonstrated that the mean BOLD response in the PPA was significantly lower for older relative to young adults when viewing scenes (i.e., the preferred stimulus type), *t*(89.34) = 4.51, *p* < 10^−4^. No age differences were present in the PPA during object trials (i.e., the non-preferred stimulus type), *t*(89.34) = .62, *p* = .535, nor were age differences present in the LOC for either objects, *t*(89.34) = 1.72, *p* = .088, or scenes, *t*(89.34) = 1.14, *p* = .257.

#### Relationship with Memory Performance

The zero-order correlations between item and source memory (averaged across image type), the PPA and LOC differentiation indices, visual acuity, and age group are shown in Table 6. Our primary hypothesis concerned the relationship between memory performance and the differentiation indices. As can be seen in Table 6, the differentiation index from the PPA, but not the LOC, was correlated with both item and source memory. Given the lack of significant correlations with the LOC, the results reported below focus solely on the PPA.

**Table 6.** Zero-order correlations between memory performance, differentiation index, similarity index, visual acuity, and age.

|  | (1) | (2) | (3) | (4) | (5) | (6) | (7) | (8) |
| --- | --- | --- | --- | --- | --- | --- | --- | --- |
| (1) Item Recognition | - |  |  |  |  |  |  |  |
| (2) Source memory | .71<br>($< .001$) | - | | | | | | |
| (3) PPA Differentiation Index | .53<br>($< .001$) | .37<br>(.010) | - | | | | | |
| (4) LOC Differentiation Index | .09<br>(.559) | .03<br>(.837) | .00<br>(.988) | - |  |  |  |  |
| (5) PPA Similarity Index | 0.50<br>($< .001$) | 0.32<br>(.026) | .78<br>($< .001$) | -.08<br>(.580) | - | | | |
| (6) LOC Similarity Index | .25<br>(.083) | .15<br>(.298) | .31<br>(.030) | .71<br>($< .001$) | .19<br>(.188) | - | | |
| (7) Visual Acuity | -.35<br>(.016) | -.16<br>(.268) | -.48<br>(.001) | .04<br>(.799) | -.45<br>(.001) | -.12<br>(.407) | - |  |
| (8) Age Group | -.28<br>(.055) | -.13<br>(.372) | -.61<br>($< .001$) | .07<br>(.632) | -.71<br>($< .001$) | -.24<br>(.106) | .63<br>($< .001$) | - |

*Note.* Correlations were computed using Pearson's $r$ . Item and source memory correlations are based on the measures after averaging across image type.

First, we focus on the correlation between item recognition and the PPA differentiation index. Importantly, this correlation remained significant after partialling out age group, *r_partial_*(45) = .48, *p* < .001 (see Figure 4A). This result, in conjunction with the absence of a moderating effect of age (see Differentiation Index Analysis in the Methods), suggests that the correlation between item recognition and the PPA differentiation index is *age invariant*. It is possible that the correlation between item recognition and PPA differentiation index is due to shared variance with source memory. Critically, the partial correlation between item recognition and PPA differentiation index controlling for both age group and source memory remained significant, *r_partial_*(44) = .33, *p* = .023 (see Figure 4B), suggesting that source memory does not account for the relationship between the differentiation index and item recognition. We also examined whether the correlation between item recognition and the PPA differentiation index was due to shared variance with visual acuity. Echoing the above analysis, the partial correlation between item recognition and the PPA differentiation index after controlling for both age group and visual acuity remained significant, *r_partial_*(44) = .46, *p* = .001.

A similar set of partial correlations to that described above was computed for the relationship between source memory performance and the PPA differentiation index. As with item recognition, the partial correlation between source memory and the PPA differentiation index was significant after controlling for age group, *r_partial_*(45) = .36, *p* = .011 (Figure 4C), and for both age group and visual acuity, *r_partial_*(44) = .35 *p* = .016. However, the correlation was no longer significant and, indeed, near zero after controlling for age group and item recognition performance, *r_partial_*(44) = .04, *p* = .779.

**Figure 4.**
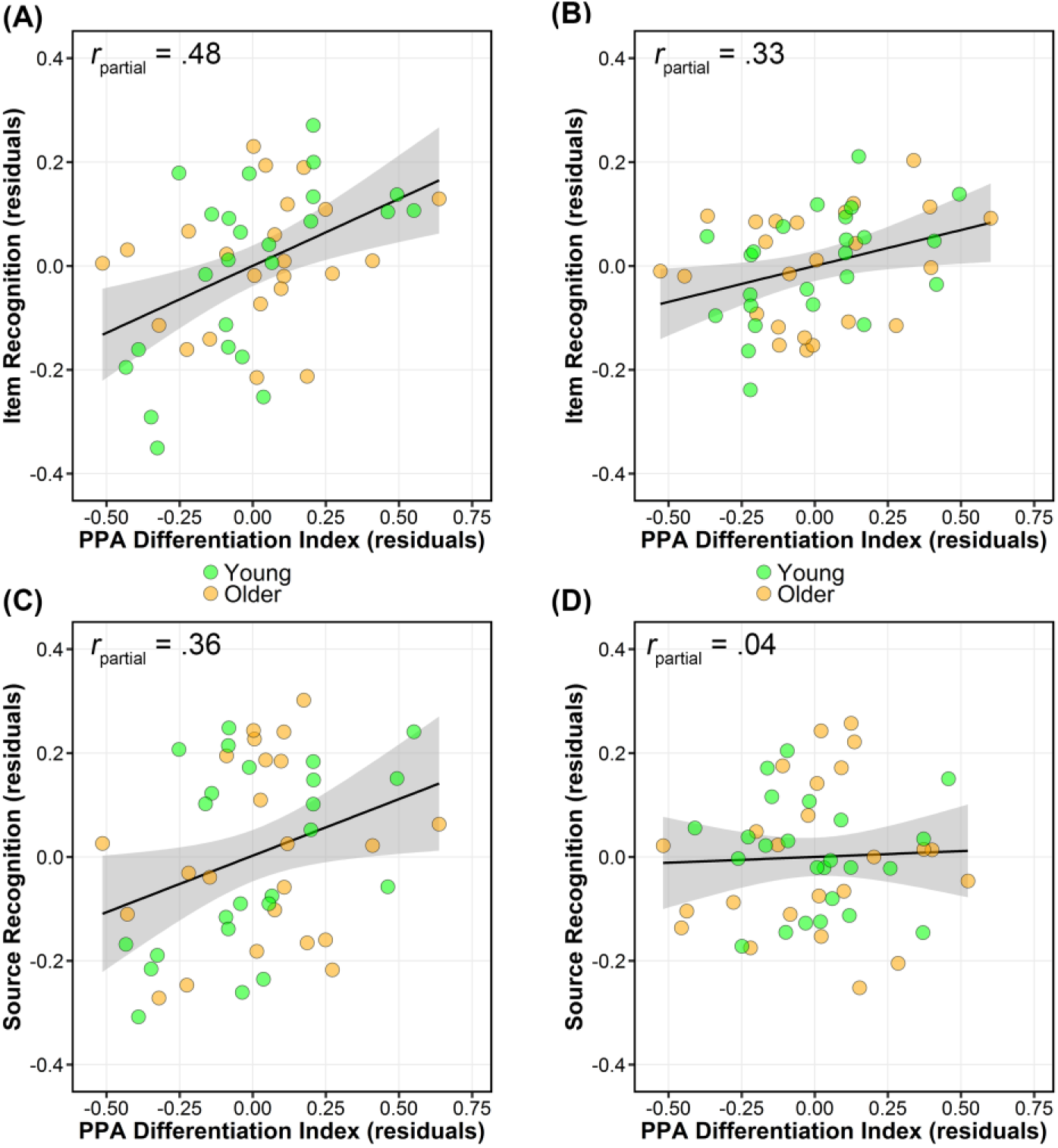
Scatter plots showing the partial correlation between the PPA differentiation index and item recognition (A,B) and source memory (C,D). The partial plots control for age group (A,C), age group and source memory (C), and age group and item recognition (D).

In summary, we observed a significant correlation between item recognition and PPA differentiation index that was *invariant* across age group, source memory performance, and visual acuity. Although the PPA differentiation index was significantly correlated with source memory, this association appeared to result from shared variance with item recognition.

#### Relationship with Neuropsychological Test Performance

Table 7 shows the zero-order correlation between the 4 neuropsychological factors (RCs), visual acuity, differentiation indices, and age group. The PPA, but not the LOC, differentiation index correlated significantly with the RCs corresponding to speed, memory, and fluency. To examine whether these correlations were independent of age, we computed partial correlations between the PPA differentiation index and the four RCs controlling for age. [It is important to reiterate that there was no indication of an interaction between age group and PPA differentiation index for any of the four RCs (see Analysis of Relationships Between Neural Differentiation and Cognition)]. The partial correlation for the speed, *r_partial_*(45) = -.09, *p* = .561, memory, *r_partial_*(45) = -.05, *p* = .759, and crystallized intelligence, *r_partial_*(45) = .11, *p* = .468, factors all failed to reach our significance threshold. Thus, the zero-order correlations between neural differentiation with the speed and memory factors reflect variance that is also shared with age group. In contrast, the partial correlation between the PPA differentiation index and the fluency factor remained significant, *r_partial_*(45) = .35, *p* = .017 (see Figure 5), suggesting that neural differentiation and fluency have an age invariant relationship. This correlation remained significant after controlling for visual acuity in addition to age, *r_partial_*(44) = .36, *p* = .014.

**Table 7.** Zero-order correlations between factor scores from the neuropsychological test performance, differentiation index, similarity index, visual acuity, and age.

|  | (1) | (2) | (3) | (4) |
| --- | --- | --- | --- | --- |
| (1) Speed (RC1) | - |  |  |  |
| (2) Memory (RC2) | -.46<br>(.001) | - |  |  |
| (3) Crystallized IQ (RC3) | .16<br>(.279) | .10<br>(.498) | - |  |
| (4) Fluency (RC4) | -.27<br>(.061) | -.27<br>(.061) | .16<br>(.287) | - |
| Correlations with: |  |  |  |  |
| PPA Differentiation Index | -.40<br>(.004) | .31<br>(.030) | .06<br>(.700) | .37<br>(.009) |
| LOC Differentiation Index | -.02<br>(.908) | .00<br>(.989) | -.08<br>(.612) | -.08<br>(.584) |
| PPA Similarity Index | -.48<br>(.001) | .34<br>(.019) | .09<br>(.560) | .30<br>(.04) |
| LOC Similarity Index | -.23<br>(.114) | .14<br>(.345) | -.05<br>(.739) | .20<br>(.182) |
| Visual Acuity | .40<br>(.005) | -.41<br>(.003) | .05<br>(.738) | -.06<br>(.677) |
| Age Group | .57<br>($< .001$) | -.56<br>($< .001$) | .05<br>(.751) | -.17<br>(.257) |

*Note.* The correlations between Visual Acuity, PPA Differentiation/Similarity Index, LOC Differentiation/Similarity Index, and Age Group are identical to those in reported in Table 6.

**Figure 5.**
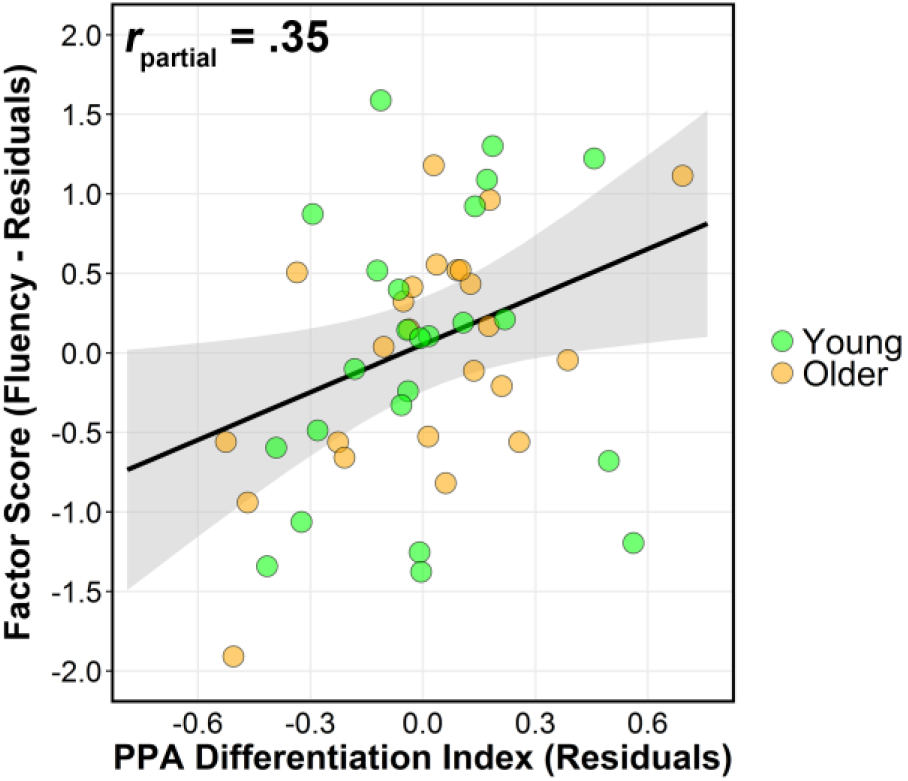
Scatter plots showing the partial correlation between the PPA differentiation index and the factor score for fluency (RC4) controlling for age.

### Pattern Similarity Index

A 2 (Age Group) X 2 (ROI) mixed ANOVA produced a significant interaction, *F*(1,46) = 25.11, *MS*_e_ = .003, *p* < 10^−5^, partial-*η*^2^ = .35 (see Figure 6a). The interaction was driven by older adults showing lower similarity indices relative to younger adults in the PPA, *t*(91.97) = 8.55, *p* < 10^−12^, but not in the LOC, *t*(91.97) = 1.40, *p* = .164. These findings mirror those observed for the univariate differentiation index and offer strong convergent evidence for age-related neural dedifferentiation in the PPA.

#### Relationship with Memory Performance

The zero-order correlations between item and source memory (averaged across image type) and the pattern similarity indices are shown in Table 6. As with the differentiation index, there were no significant correlations involving the LOC similarity index. Thus, we focus the partial correlation analysis on the index from the PPA. The correlation between item recognition and the PPA similarity index remained significant after partialling out age group, *r_partial_*(45) = .45, *p* = .002 (see Figure 6B). This result, in conjunction with the absence of a moderating effect of age (see Pattern Similarity Analysis in the Methods), suggests that the correlation between item recognition and the similarity index in the PPA is *age invariant*. Moreover, the correlation remained significant after partialling out both age group and source memory performance, *r_partial_*(44) = .33, *p* = .025, and age group and visual acuity, *r_partial_*(44) = .46, *p* = .002. These latter two results suggest that the correlation between item recognition and the PPA similarity index was not driven by variance shared with source memory or visual acuity, respectively.

The correlation between source memory and the PPA similarity index was also age invariant, *r_partial_*(45) = .32, *p* = .026. Although this correlation remained significant when partialling out age group and visual acuity, *r_partial_*(44) = .32, *p* = .028, adding item recognition as a covariate along with age group rendered the correlation non-significant, *r_partial_*(45) = .01, *p* = .946. Thus, the results using the pattern similarity index parallel those for the differentiation index in that the metric of neural differentiation predicted item, but not source, memory in an age invariant manner.

**Figure 6.**
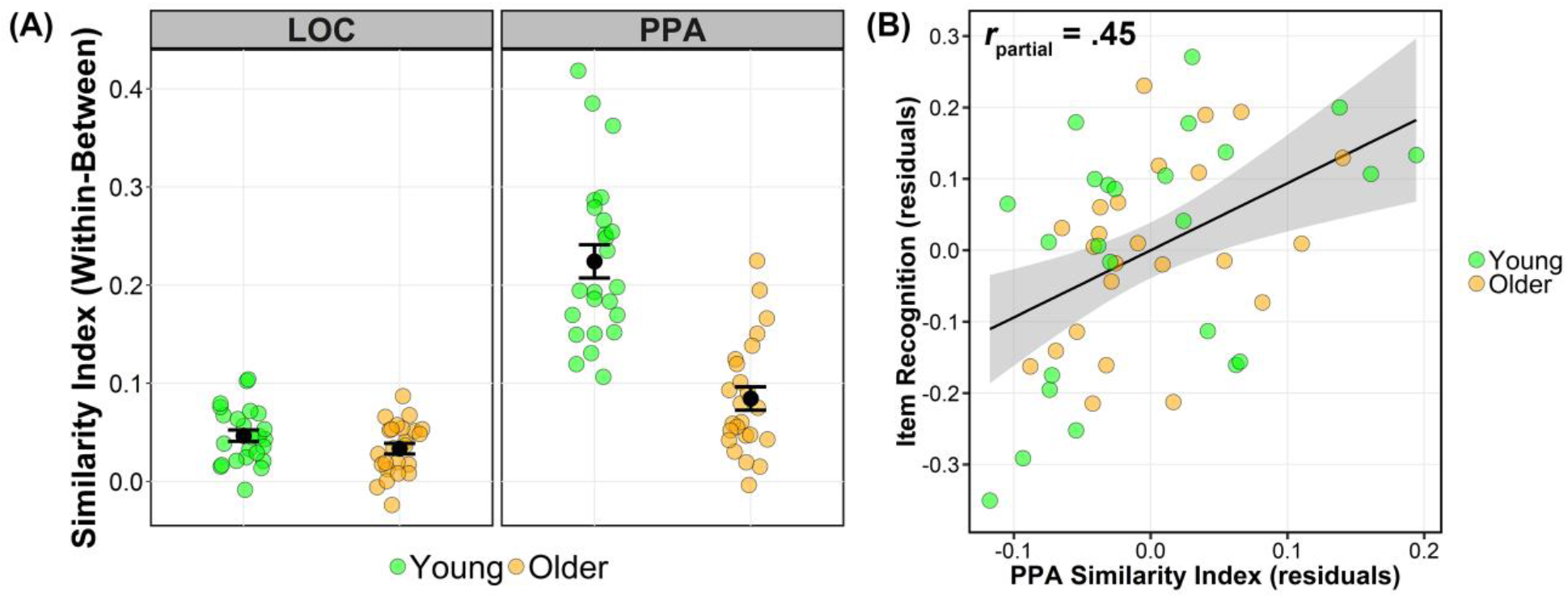
(A) Plot of the similarity index (within-between similarity for the preferred image type) computed from the LOC and PPA for young and older adults. (B) Scatter plot showing the partial correlation between the similarity index in the PPA and item recognition controlling for age group.

#### Relationship with Neuropsychological Test Performance

Table 7 shows the zero-order correlation between the 4 neuropsychological factors (RCs) and the PPA and LOC similarity indices. Again, we focus on the PPA as none of the zero-order correlations for the LOC similarity index reached our significance threshold. The partial correlation for the speed, *r_partial_*(45) = -.13, *p* = .367, memory, *r_partial_*(45) = -.10, *p* = .512, crystallized intelligence, *r_partial_*(45) = .17, *p* = .258, and fluency, *r_partial_*(45) = .26, *p* = .080, factors all failed to reach our significance threshold after controlling for age group. The lack of a significant partial correlation between the PPA similarity index (controlling for age group) and the fluency factor stands in contrast to findings for the differentiation index reported above. It is noteworthy, however, that the correlation was sizeable and in the same direction as that for the differentiation index.

## Discussion

We describe three main findings. First, we replicated prior findings (e.g., Park et al., 2004, 2012;Voss et al., 2008) by showing age-related reductions in two measures of category-level neural differentiation (henceforth, collectively termed neural differentiation indices). These age differences were observed only in the PPA, and not in the LOC. Second, we found an age invariant relationship between neural differentiation in the PPA and item recognition memory. Lastly, a similarly age invariant relationship was evident between a ‘fluency’ factor derived from neuropsychological test scores and neural differentiation (albeit, reaching significance only for the differentiation index). Together, the findings suggest that neural differentiation in the PPA is associated with two independent sources of variance: age and cognitive performance.

### Absence of Age Differences in Item and Source Memory

No age differences were observed in study RT, item recognition, or source memory. While age differences in RT might be expected, null age effects on study RT have been reported previously in tasks very similar to the present one (e.g., de Chastelaine et al., 2011, 2016;Mattson et al., 2014; Wang et al., 2016). The lack of an age difference in source memory is more surprising given well-documented age-related deficits in recollection (Koen and Yonelinas, 2014; Schoemaker et al., 2014) and source memory (Spencer and Raz, 1995; Old and Naveh-Benjamin, 2008). This null finding might reflect our employment of an atypical older sample. This is a perennial concern in neuroimaging studies of aging (Rugg, 2017), but is mitigated here by the ‘standard’ pattern of impaired and preserved neuropsychological test performance demonstrated by our older participants (e.g., Drag and Bieliauskas, 2010; Park et al., 2002). A second possibility is that age differences in source memory were masked by an especially conservative response bias in young adults. This could have resulted from our instruction to report source memory decisions only when confidence was high. In complying, young adults might have withheld what would have been accurate decisions because their response criteria were set above the threshold necessary for accurate responding, lowering their source accuracy and attenuating potential age differences. Lastly, the encoding tasks might have disproportionately benefited memory encoding in older adults, an effect that has sometimes been reported to eliminate age differences in recollection (Luo et al., 2007). Although the last two accounts are not mutually exclusive, the latter account also accommodates the null age effects on item memory.

### The Age Component of Neural Differentiation

Our findings replicate prior research demonstrating that age-related neural dedifferentiation in the PPA is driven by diminished BOLD responses to scenes in older adults (“neural attenuation”;Park et al., 2012). Counter to prior findings (Park et al., 2004; for related findings, see Berron et al., 2018), we did not observe significant age differences in neural differentiation in the LOC, a region selectively responsive to objects from a wide variety of categories (Grill-Spector et al., 2001). This null finding for the LOC is not unprecedented: Chee and colleagues (2006) also reported null age differences in the LOC for objects (relative to scenes); relatedly, Voss and colleagues (2008) reported null effects of age on neural selectivity for familiar words and colors.

Our results add to the evidence for age-related neural dedifferentiation, but do little to elucidate its functional significance. Any account must, however, accommodate the present and prior findings (see above) that age-related dedifferentiation is evident only for some stimulus classes. One possibility (raised by a reviewer) is that the present findings have their origin not in the way different neural regions represent visual categories as a function of age, but in age-related differences in eye-movements. By this argument, the results for the PPA reflect the adoption by older and younger adults of different scanning strategies when confronted with scenes (e.g., Açık et al., 2010). This account cannot be definitively ruled out in the absence of eye-movement data (which, to our knowledge, have yet to be reported in any relevant study). We note however that it cannot be a general explanation of age-related neural differentiation, which has been reported not only for visual stimuli, but for auditory stimuli and motoric activity also (Carp et al., 2011a; Grady et al., 2011a, 2011b).

A second account arises from the prosaic idea that perceptual experience and knowledge accumulate over the lifespan because of an ever-increasing number of encounters with new exemplars of different perceptual categories (for related findings showing that the neural correlates of object processing are moderated by a variable related to life experience, namely culture, see Goh et al., 2007; for review, see Goh and Park, 2009). Thus, when confronted with a novel exemplar, older individuals are arguably better able to assimilate it into a pre-existing representational structure (a perceptual “schema”;Gilboa and Marlatte, 2017) than are young adults, who have had less opportunity to develop such schemas. Consequently, with increasing age, perceptual processing of novel category exemplars will come to more closely resemble the processing afforded previously experienced exemplars. By this hypothesis, therefore, age-related neural dedifferentiation is not necessarily a detrimental consequence of increasing age.

This ‘familiarity hypothesis’ accounts for two important aspects of the present data. First, it is consistent with the findings that age-related dedifferentiation in the PPA resulted from neural attenuation. According to the above hypothesis, the processing of novel exemplars of a visual category will more closely resemble the processing engaged by familiar exemplars in older than in younger adults. Thus, when first encountered, such stimuli might be expected to elicit smaller neural responses in older individuals, that is, to demonstrate ‘repetition suppression’ – the much-studied neural correlate of perceptual priming (e.g., Henson and Rugg, 2003; Gotts et al., 2012; Barron et al., 2016).

Second, the hypothesis provides an explanation for the absence of age-related neural dedifferentiation in the LOC reported here and previously (Chee et al., 2006), and its absence in word- and color-selective cortical regions in Voss and colleagues (2008). The hypothesis predicts that age differences in neural differentiation will be diminished for exemplars that are similarly familiar to both young and older individuals. Arguably, even young adults have experienced canonical objects of the kinds employed in the present study on numerous occasions prior to the experimental session, resulting in a blunting of age-differences in neural differentiation. Consistent with this proposal, Voss and colleagues (2008) failed to identify age-related dedifferentiation for words, whereas Park and colleagues (2004) reported robust dedifferentiation for pseudo-words, items that likely would not have been encountered by members of either age group pre-experimentally.

### Relationship Between Neural Differentiation and Memory Performance

We observed robust correlations between the PPA neural differentiation index and both recognition memory performance for the experimental items, and a fluency factor derived from neuropsychological test scores (for related findings, see Park et al., 2010; Du et al., 2016; Berron et al., 2018). The finding that lower neural differentiation was predictive of poorer memory performance is broadly consistent with our pre-experimental hypothesis that dedifferentiation should impact memory encoding. Importantly, this relationship was age invariant, and suggests that neural selectivity and item recognition are similarly coupled across much of the adult lifespan (Rugg, 2017). As suggested by a reviewer, our failure to find age differences in memory performance might have contributed to the failure to find a moderating effect of age on the relationships between neural differentiation and cognitive performance. While we cannot definitively rule out this possibility, we note that findings from prior studies indicate that null effects of age on a behavioral measure are not a precondition for finding age-invariant brain-behavior correlations (e.g., de Chastelaine et al., 2011, 2016;Wang et al., 2016; for related findings, see Du et al., 2016).

Another important result is the seemingly selective relationship between neural differentiation and item recognition. Whereas the correlation with recognition remained when source memory performance was controlled for, the reverse was not the case. Thus, neural differentiation was primarily a predictor of memory for the experimental items themselves, and not for their study contexts, possibly suggesting that the relationship between neural differentiation and memory performance is dependent on such factors as task demands. One might predict that a unique relationship between source memory performance and neural differentiation would have emerged had the studied scenes and objects been employed as source features rather as test items.

As noted, we found an age invariant relationship between neural differentiation and one of the latent factors – ‘fluency’ – derived from neuropsychological test performance. In line with Park and colleagues (2010), who described an analogous relationship between neural differentiation and fluid intelligence (in older adults only), the present finding suggests that neural differentiation may index not just the precision with which perceptual information is represented, but also broader aspects of neural efficiency. More generally, our findings that the relationships between neural differentiation and item memory performance and fluency were age invariant could be seen as a challenge to the view that neural dedifferentiation is a determinant of cognitive aging (e.g., Li et al., 2001; Park et al., 2010). This conclusion should be treated as provisional, however, until the present findings are replicated in larger and more diverse samples of participants.

